# Siderophore production and utilization by microbes in the North Pacific Ocean

**DOI:** 10.1101/2022.02.26.482025

**Authors:** Jiwoon Park, Bryndan P. Durham, Rebecca S. Key, Ryan D. Groussman, Paulina Pinedo-Gonzalez, Nicholas J. Hawco, Seth G. John, Michael C.G. Carlson, Debbie Lindell, Laurie Juranek, Sara Ferrón, Francois Ribalet, E. Virginia Armbrust, Anitra E. Ingalls, Randelle M. Bundy

## Abstract

Siderophores are strong iron-binding molecules produced and utilized by microbes to acquire the limiting nutrient iron (Fe) from their surroundings. Despite their importance as a component of the iron-binding ligand pool in seawater, data on the distribution of siderophores and the microbes that use them are limited. Here we measured the concentrations and types of dissolved siderophores during two cruises in April 2016 and June 2017 that transited from the iron-replete, low-macronutrient North Pacific Subtropical Gyre (NPSG) through the North Pacific Transition Zone (NPTZ) to the iron-deplete, high-macronutrient North Pacific Subarctic Frontal Zone (SAFZ). Surface siderophore concentrations in 2017 were higher in the NPTZ (4.0 – 13.9 pM) than the SAFZ (1.2 – 5.1 pM), which may be partly attributed to stimulated siderophore production by environmental factors such as dust-derived iron concentrations (up to 0.51 nM). Multiple types of siderophores were identified on both cruises, including ferrioxamines, amphibactins and iron-free forms of photoreactive siderophores, which suggest active production and use of diverse siderophores across latitude and depth. Additionally, the widespread genetic potential for siderophore biosynthesis and uptake across latitude and the variability in the taxonomic composition of bacterial communities that transcribe putative ferrioxamine, amphibactin and salmochelin transporter genes at different latitudes suggest that particular microbes actively produce and use siderophores, altering siderophore distributions and the bioavailability of iron across the North Pacific.

## Introduction

Iron (Fe) is an essential nutrient for ocean life, but due to its low solubility in seawater, it often exists in concentrations far below the levels required for sustaining phytoplankton growth in over a third of open ocean surface waters (Johnson et al. 1997; Liu and Millero 2002; Moore et al. 2013). More than 99% of the dissolved iron in the ocean is bound to organic ligands, as complexation of iron to organic ligands can increase the solubility of iron by orders of magnitude and prevent precipitation and scavenging (Gledhill and van den Berg 1994; Rue and Bruland 1995; Hunter and Boyd 2007; Gledhill and Buck 2012). In addition to increasing iron solubility, these organic ligands also affect the bioavailability of iron to phytoplankton and bacteria, playing a critical role in the ocean iron and other biogeochemical cycles (Hassler et al. 2012; Shaked et al. 2020).

Although the importance of iron-binding ligands to iron speciation and distributions in the ocean has been well recognized, the identity of these ligands, their sources and the conditions in which they are produced remain unclear (Tagliabue et al. 2017). Iron-binding ligands in seawater have been most commonly measured by competitive ligand exchange-adsorptive cathodic stripping voltammetry (CLE-ACSV) (Gledhill and van den Berg 1994; Rue and Bruland 1995; Buck et al. 2012; Bundy et al. 2015). This electrochemical method groups ligands into classes operationally defined by their binding affinity for iron, with the L_1_ class being the strongest iron-binding ligands and L_2_ and below being progressively weaker ligands (Hunter and Boyd 2007; Gledhill and Buck 2012; Bundy et al. 2014, 2015). As the concentration of the L_1_ ligand class is closely correlated to dissolved iron throughout the global ocean (Gledhill and Buck 2012; Buck et al. 2015, 2018), it is currently the main class of ligands represented in biogeochemical models of the iron cycle (Tagliabue et al. 2017). CLE-ACSV does not provide any information on the chemical structures of different ligands across classes, but combining these method with mass spectrometry allows direct characterization of individual ligand structures at the molecular level (Mawji et al. 2008; Boiteau et al. 2013, 2016; Velasquez et al. 2016; Bundy et al. 2018). The similarities between the iron binding strength of L_1_ class ligands and that of siderophores suggest that a part of the L_1_ ligand pool may be made up of siderophores (Witter et al. 2000; Macrellis et al. 2001; Vraspir and Butler 2009).

Siderophores are low molecular weight, strong iron-binding molecules produced by microbes (Neilands 1981). More than 500 siderophores have been identified and isolated to date, mostly from bacteria and fungi cultures (Sandy and Butler 2009; Hider and Kong 2010), Siderophores are synthesized inside cells by either the non-ribosomal peptide synthetase (NRPS) pathway or the NRPS-independent siderophore (NIS) synthetase pathway and are secreted to the environment (Gulick 2017; Carroll and Moore 2018). Gram-negative bacteria use a combination of TonB-dependent outer membrane receptors (TBDT) and ATP-binding cassette (ABC) transporters to transport iron-bound siderophores through the outer membrane into the periplasm and the cytoplasm, where iron gets reduced from Fe(III) to Fe(II) and released from siderophores (Krewulak and Vogel 2008; Schalk and Guillon 2013). The ability to synthesize and transport siderophores varies widely between microbes: some microbes can produce one or more siderophores, while some cannot produce siderophores themselves but possess the siderophore uptake machinery to use exogenous siderophore-bound iron, and some cannot produce or utilize siderophores (Sandy and Butler 2009; Gledhill and Buck 2012). Siderophore production benefits its producers by decreasing the amount of available iron to other microbes that lack the ability to acquire siderophore-bound iron. In addition, as siderophores can solubilize particulate iron to increase the amount of dissolved iron, they can alter the pool of bioavailable iron in the marine environment and shape the microbial community structure and composition (Kraemer et al. 2005; Kügler et al. 2020).

Multiple studies to date have measured picomolar concentrations of siderophores in surface and mesopelagic ocean waters (0 – 1,500 m) (Mawji et al. 2008; Velasquez et al. 2016; Boiteau et al. 2016, 2019; Bundy et al. 2018). In these studies, characterized iron-siderophore complexes accounted for 0.2 - 10% of the total dissolved iron pool, which suggests that siderophores can play a significant role in ocean iron cycling. Indeed, the prevalence of siderophore-based iron acquisition in the ocean has been demonstrated by genomic and transcriptomic surveys of marine communities. For instance, TonB-dependent outer membrane receptors involved in siderophore uptake were widespread among prokaryotes in the Global Ocean Survey (GOS) metagenomes, and prokaryotes in the Southern Ocean preferred the siderophore-mediated iron uptake pathway to ferrous and ferric iron transport pathways (Hopkinson and Barbeau 2012; Toulza et al. 2012; Tang et al. 2012; Debeljak et al. 2019). Siderophore biosynthesis genes were detected less frequently and at a lower abundance from these genomes than siderophore uptake genes, potentially because fewer siderophore synthesis genes were characterized and included in the search databases, or because most microbes cannot afford the expensive cost of siderophore production (Hopkinson and Barbeau 2012; Toulza et al. 2012). While metagenomes give insights into the potential for biosynthesis or uptake of siderophores, transcription of siderophore biosynthesis and uptake genes in environmental transcriptomes can assess active production and utilization of siderophores in the ocean and provide evidence for possible rapid cycling of these compounds in seawater. However, to our knowledge, no study to date has paired a genomics- and transcriptomics-based approach with chemical measurements of siderophore concentrations and distributions.

To examine possible chemical and biological controls of siderophore distributions, we measured dissolved siderophore concentrations across a basin-scale meridional transect at 158°W, from 23.5°N to 41.4°N in the North Pacific Ocean during two cruises in 2016 and 2017. In order to evaluate the impact of microbial activity on siderophore distribution, we searched for a comprehensive set of siderophore synthesis and uptake genes across both cruise transects, then further looked at transcription of transporter genes for three specific siderophores.

## Methods

### Seawater Collection

Samples were collected during the Gradients 1 cruise (KOK1606) on the R/V *Ka’imikai-O-Kanaloa* from April 19 to May 3, 2016 and the Gradients 2 cruise (MGL1704) on the R/V *Marcus G. Langseth* from May 27 to June 13, 2017. The two cruises collected samples along 158°W, from approximately 23.5°N to 37.3°N and 25.8°N to 41.4°N respectively. Seawater samples for dissolved siderophore analyses were collected using Niskin-X bottles with external Teflon-coated springs (General Oceanics) mounted on a trace metal-clean rosette and a non-metallic line. Collection depths were pre-programmed to the autofire module (Seabird Scientific) and sample bottles were closed on the upcast while the rosette was moving through the water (∼ 20-30 m min^-1^). Niskin-X bottles were placed in a HEPA-filtered clean space on board the ship prior to sub-sampling. Surface samples (∼ 5 m) were collected from a towed trace metal-clean “fish” (Vink et al. 2000) and sample bottles were filled inside a HEPA-filtered clean space.

Sea surface temperature and salinity were measured from thermosalinograph systems aboard each research vessel. Sample collection and analysis for nitrate plus nitrite (N+N), soluble reactive phosphorus (SRP), particulate organic carbon (POC) concentrations and O_2_/Ar-derived estimates of net community production (NCP) are described in Juranek et al. (2020).

Discrete samples for cell abundance measurements were collected from surface water (∼ 5 m depth), fixed with glutaraldehyde (0.25% final concentration) and analyzed with a BD Influx cell sorter equipped with a 488 nm laser. Data collection was triggered by forward light scatter (10% neutral density). Total bacteria counts were obtained by staining the sample with SYBR Green I (0.01% final concentrations) for 20 minutes in the dark and detecting emission at 530 nm (40 nm bandpass). *Prochlorococcus* cell counts were determined on unstained samples based on forward scatter and chlorophyll fluorescence emission at 692 nm (40 nm bandpass). Heterotrophic bacteria abundance was calculated by subtracting *Prochlorococcus* counts from the total bacteria count (Marie et al. 1999).

### Metagenomic and metatranscriptomic sampling and processing

During both cruises, seawater samples for metagenomes and metatranscriptomes were collected either from the flow-through system or from a CTD rosette at 15 m depth. In each instance, 6-10 liters of seawater were passed through a nitex pre-filter (200 μm mesh for Gradients 1 and 100 μm mesh for Gradients 2) followed by sequential filtration through a 3 μm and a 0.2 μm polycarbonate filters using a peristaltic pump. This collection scheme yielded two size classes (0.2 – 3 µm and 3 – 100 or 200 µm). In this study, we focused on the 0.2 – 3 µm size fraction that includes “free-living” bacterial communities. Filters were frozen in liquid nitrogen and stored at −80°C until further processing.

Samples from the 0.2 – 3 µm size fraction were used to construct sequencing libraries. To generate quantitative read inventories, internal standards of *T. thermus* genomic DNA for metagenomes and a set of 14 synthetic internal mRNA standards for metatranscriptomes were added in known concentrations prior to nucleic acid extraction. DNA was extracted using phenol:chloroform following enzymatic cell lysis (Boström et al. 2004). DNA was fragmented to ∼ 600 bp and used to construct Nextera DNA Flex libraries. Total RNA was extracted using the ToTALLY RNA kit (Invitrogen) for Gradients 1 samples and the Direct-zol RNA MiniPrep kit (Zymo Research) for Gradients 2 samples, and processed as described previously (Durham et al. 2019). Briefly, RNA extracts were treated with DNase, and rRNAs were removed using Illumina’s Bacteria and Yeast Ribo-Zero rRNA Removal Kits. The rRNA-depleted RNA was cleaned using the Zymo RNA Clean and Concentrator Kit. Purified, depleted RNA was then sheared to ∼ 225 bp fragments and used to construct TruSeq cDNA libraries according to the Illumina TruSeq® RNA Sample Preparation v2 Guide. Metagenomic and metatranscriptomic libraries were sequenced with the Illumina NovaSeq 6000 sequencing platform with paired-end (2 × 150) chemistry.

### Dissolved iron analyses

Surface seawater samples for dissolved trace metals were filtered through 0.2 µm Supor membrane filters (Pall), and the filtrates were stored in 50 mL acid-cleaned polypropylene centrifuge tubes (VWR). Samples were analyzed after off-line preconcentration onto Nobias PA1 chelating resin at pH ∼ 6.5 using the seaFAST-pico system (Lagerström et al. 2013). Samples were eluted in 10% nitric acid (Optima, Fisher Scientific) and analyzed on a Thermo Fisher Element 2 High Resolution ICP-MS using the isotope dilution method (Pinedo-González et al. 2020). The accuracy of measured dissolved iron concentrations was evaluated by referencing to consensus values of a seawater reference standard (GEOTRACES GS) and are reported elsewhere (Gradoville et al. 2020; Pinedo-González et al. 2020).

### Dissolved siderophore sample collection and analysis

Ten to twenty liters of 0.2 µm filtered (Acropak 200; Pall Corporation) seawater were collected from stations between 23.5°N and 37.5°N (Gradients 1) or between 29.5°N and 41.4°N (Gradients 2) from multiple depths between the surface and 400 m. The filtered seawater was pumped continuously at a flow rate of 18 mL/min through a Bond Elut column (1 g ENV, 6 mL, Agilent Technologies) that had been previously cleaned with two column volumes of pH 2 Milli-Q water and two column volumes of Milli-Q water, and activated with two column volumes of trace metal clean methanol (Fisher Scientific Optima or distilled methanol). After the sample solid phase extraction was completed, columns were washed with 12 mL of Milli-Q water and stored at −20°C for later analyses.

Prior to analysis in the laboratory, the Bond Elut columns were thawed, rinsed with two column volumes of Milli-Q water and eluted using 13 mL of optima or distilled methanol. The eluents were dried down using a vacuum concentrator with a refrigerated vapor trap (Thermo Scientific) over 4 to 5 hours, brought up to final volumes of 500 µL using Milli-Q water and stored in 2 mL low density polyethylene vials. Multiple method blanks were prepared along with samples to ensure that the samples were not contaminated during extraction. All samples were analyzed using liquid chromatography coupled to inductively coupled plasma mass spectrometry (LC-ICP-MS) and to electrospray ionization mass spectrometry (LC-ESI-MS). For each analysis, 50 µ L aliquots of each sample were spiked with 10 µ L of 50 µM cyanocobalamin as an internal standard and were analyzed on a Dionex Ultimate 3000 liquid chromatography system with a polyetheretherketone (PEEK) ZORBAX-SB C18 column (4.2 × 150 mm, 3 µm, Agilent Technologies). Samples were separated using a flow rate of 50 µ L/min at 30°C and a 20 minute gradient from 95% solvent A (Milli-Q water with 5 mM ammonium formate) and 5% solvent B (trace metal clean methanol with 5 mM ammonium formate) to 90% solvent B, followed by a 10 minute isocratic step of 90% solvent B, then a 5 minute gradient from 90% to 95% solvent B, an isocratic 5 minute step at 95% solvent B, and a 15 minute conditioning step at 5% B prior to injecting the next sample. The same chromatography scheme was used both for the LC-ICP-MS and the LC-ESI-MS analyses (Boiteau et al. 2016; Bundy et al. 2018).

Samples were introduced from the LC to the ICP-MS (iCAP-RQ; Thermo Scientific) equipped with platinum sample and skimmer cones at a flow rate of 50 µ L/min via a PFA-ST nebulizer (Elemental Scientific) and a spray chamber at 2.7°C. A 10% oxygen flow was added to the sample gas to prevent deposition of organic materials on the sampler and skimmer cones. Measurements were made in kinetic energy discrimination (KED) mode with a helium collision gas at a flow rate of 3.8 – 4.0 mL/min. ^56^Fe peaks were identified using in-house R scripts (Boiteau et al. 2016; Bundy et al. 2018), and siderophore concentrations were calculated from calibration curves of ferrioxamine E (Boiteau et al. 2016; Bundy et al. 2018).

To identify siderophores structurally, samples were further analyzed with an Orbitrap (Q-Exactive HF; Thermo Scientific) with the following instrument settings: 3.5 kV spray voltage, 320°C capillary temperature, 16 sheath gas, 3 auxiliary gas and 1 sweep gas (arbitrary units), 90°C auxiliary gas heater temperature and S-lens RF level 65. MS^1^ scans were collected in full positive mode with 120,000 mass resolution, 200 – 2000 m/z mass range, 1e^6^ AGC target, and 100 ms maximum injection time. MS^2^ scans were collected in data-dependent MS^2^ (dd-MS^2^) mode with 30,000 mass resolution, 1.0 m/z isolation window, 2e^4^ AGC target, 100 ms maximum injection time and 35% collision energy, and used an inclusion list containing known masses of ∼ 300 siderophores. Raw data files were converted to mzXML formats using MSConvert (Proteowizard) and processed via in house R scripts using the XCMS package (Boiteau et al. 2016; Bundy et al. 2018). Cyanocobalamin peaks (m/z = 678.234) in each extracted ion chromatogram were aligned to ^59^Co peaks in each ICP-MS chromatogram to account for differences in retention times. Siderophores were identified using exact masses of known siderophore compounds from MS^1^ (± 0.005Da), then further identified by MS^2^ fragmentation patterns when possible.

A confidence level was assigned to each mass feature based on the putative identification of each siderophore. The confidence levels were modelled after those used in metabolite identification (Sumner et al. 2007) (Supp. Table 1). Level 1 is the highest level of confidence, for which MS^2^ spectra of our samples were matched to spectra from other publications (10 compounds). MS^2^ spectra were not obtained on all samples, so we matched exact masses (± 0.005Da) and retention time (± 0.5 minutes) from samples with MS^2^ data to identify compounds in samples with no MS^2^ data, and annotated those compounds as level 2 (13 compounds). Compounds that had MS^2^ spectra matching in silico MS^2^ spectra (CFM-ID and MetFrag) were annotated as level 3 (4 compounds) (Ruttkies et al. 2016; Djoumbou-Feunang et al. 2019). Lastly, compounds were annotated as level 4 if they were identified as a putative siderophore based on MS^1^ but there was no MS^2^ spectra available for comparison, or if the obtained MS^2^ spectra did not agree with the in silico predicted MS^2^ spectra (61 compounds).

### Metagenome and metatranscriptomes analysis

Sequence reads from metagenomes were quality controlled using BBDukv38.22 with the settings ktrim=r, k=23, mink=8, hdist=1, qtrim=rl, trimq=20, minlen=50, tpe, maq=10. Reads were then assembled into contigs de novo using SPAdesv3.13.1 using the --meta flag with kmer lengths of 33, 55 and 77. Metagenomes were analyzed using FeGenie, a tool that identifies potential iron metabolism related genes, with FeGenie’s HMM library for siderophore biosynthesis and transport genes (Garber et al. 2020).

Raw Illumina metatranscriptome data was quality controlled with trimmomatic v0.36 (Bolger et al. 2014) using the parameters *MAXINFO:135:0*.*5, LEADING:3, TRAILING:3, MINLEN:60*, and *AVGQUAL:20*, and matching read pairs were merged using flash v1.2.11 (Magoc and Salzberg 2011) with parameters -*r 150 -f 250 -s 25*. FASTQ files were converted to FASTA and translated with seqret and transeq vEMBOSS:6.6.0.0 (Rice et al. 2000) using standard genetic code. Potential coding frames with ≥ 40 uninterrupted amino acids were retained for further analysis.

We searched for translated reads corresponding to outer membrane transporters of three siderophores – ferrioxamine (FoxA and DesA), amphibactin (FhuA), and salmochelin (FepA and IroN). Full-length, experimentally verified protein sequences from reference genomes (Supp. Table 2) were used as queries to identify putative orthologs in translated protein sequences from marine microbial genomes and transcriptome assemblies compiled in a custom marine reference database (Coesel et al. 2021) using BLASTp v2.2.31. Genomes and transcriptomes used in this database were derived from Joint Genome Institute (JGI), National Center for Biotechnology Information (NCBI), and the Marine Microbial Eukaryote Transcriptome Sequence Project (MMETSP) (Keeling et al. 2014). A complete list is available in Coesel et al. (2021). Marine sequences retrieved from the BLASTp search were clustered using usearch (Edgar 2010) and aligned using MAFFT v7 with the E-ISN-I algorithm (Katoh and Standley, 2013). The alignment was trimmed using trimAl v1.2 using *– gt .05 –resoverlap 0*.*5 –seqoverlap 50* options (Capella-Gutierrez et al. 2009). The trimmed alignment file was converted to PHYLIP format, and the best-fit amino acid substitution matrix, among-site rate heterogeneity model, and observed amino acid frequency were determined using ProtTest 3 software (Darriba et al. 2011). A maximum-likelihood phylogenetic reference tree was built using RAxML v8 (Stamatakis 2014), and only those sequences that clustered with experimentally verified enzymes were considered putative homologs. These phylogenetic reference trees served as scaffolds to recruit translated environmental metatranscriptomic reads. An HMM profile was constructed from each reference alignment using hmmbuild, followed by transcript identification and alignment to the reference using hmmsearch (parameters: *-T 40 –incT 40*) and hmmalign, respectively, with the HMMER package v2.1 (Eddy 2011). NCBI taxonomy was assigned to each environmental sequence using pplacer v1.1.alpha19-0-g807f6f3 (Matsen et al. 2010) based on the read placement with the best maximum likelihood score to the reference tree (parameters: *--keep-at-most 1 –max-pend 0*.*7*). Read counts were normalized by recovery of internal mRNA standards to estimate transcript abundance in total transcripts per liter. Due to the high similarity between ferrioxamine, amphibactin, and ferrichrome transporter sequences (48 – 100%) and between salmochelin and enterobactin transporter sequences (58 – 100%), each group of transporter homologs was combined to construct a single reference tree (Supp. Fig. 1; Supp. Table 1). For the combined ferrioxamine-amphibactin-ferrichrome transporters, we report sequences most closely related to the experimentally verified sequences separately from those more distantly related (Supp. Fig. 1; Supp. Table 1). Read counts from triplicate samples were averaged at each latitude. To determine if the mean read counts for each taxonomic group were significantly different between latitudes, a one-way ANOVA and a post hoc Tukey multiple comparison test was applied.

## Results

### Environmental characteristics of the North Pacific Transition Zone and surrounding waters

The two cruises crossed three distinct regions of the North Pacific Ocean: the North Pacific Subtropical Gyre (NPSG), the North Pacific Transition Zone (NPTZ), and the North Pacific Subarctic Frontal Zone (SAFZ). The NPTZ has previously been defined based on latitudinal gradients of physical and biological features (Roden 1991; Polovina et al. 2001). The southern boundary of the NPTZ, or the North Pacific subtropical front, generally corresponds to the 34.8 isohaline and the 18°C isotherm (Roden 1991). The transition zone chlorophyll front (TZCF) is a dynamic feature between the NPSG and the subarctic gyre, and is operationally defined by surface chlorophyll-*a* concentrations of 0.2 mg/m^3^ (Polovina et al. 2001). The TZCF does not always coincide with the physical definition of the NPTZ, as it migrates seasonally by ∼10 degrees in latitude, due to strong seasonal gradients in nutrients, light and shifts in phytoplankton composition (Follett et al. 2021).

In both cruise transects, we observed decreases in sea surface temperature and salinity as we transited northward (Fig. 1A, B). The second derivative of surface salinity was used to determine the position of the North Pacific subtropical front, which was 30.7°N in April 2016 and 32.8°N in June 2017 (Gradoville et al. 2020; Follett et al. 2021). In addition, the subarctic salinity front was only crossed in June 2017, and this front was located at 38.0°N based on the 33.8 isohaline (Roden 1991; Gradoville et al. 2020). These frontal locations will be used in the remainder of this work when discussing the NPSG, the NPTZ, and the SAFZ (Fig. 1).

**Figure 1.**
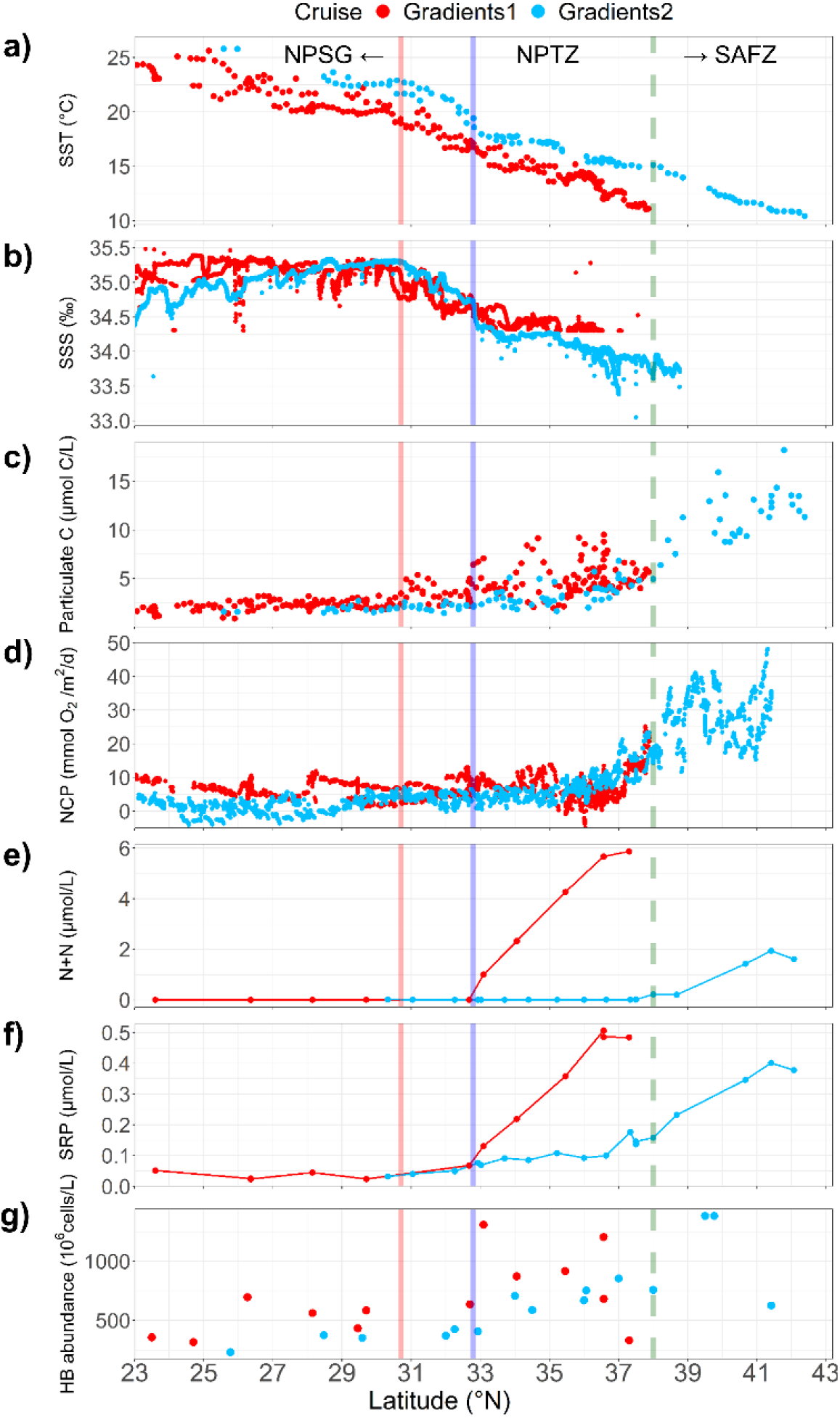
Physical, chemical and biological properties measured along the two cruise transects. A) Sea surface temperature (SST), B) sea surface salinity (SSS), C) nitrate and nitrite (N+N) concentrations, D) soluble reactive phosphorus (SRP) concentrations, E) particulate organic carbon concentrations, F) O_2_/Ar-derived estimates of net community production (NCP) and G) heterotrophic bacterial abundances from Gradients 1 (red) and 2 (blue) cruises. Vertical red and blue solid lines indicate location of subtropical fronts from Gradients 1 and 2 cruises respectively; green dashed line indicates location of the subarctic front from Gradients 2.

Strong latitudinal trends in surface nutrient inventories were observed in both cruises, with N+N and SRP concentrations both increasing across the NPSG to NPTZ or SAFZ by up to tenfold (Juranek et al. 2020) (Fig. 1C, D). Along with changes in nutrients, latitudinal changes in productivity and biomass were also observed. Both particulate organic carbon (POC) and O_2_/Ar-derived net community production (NCP) sharply increased across the TZCF, which was located at 33.0°N in April 2016 and 36.2°N in June 2017, and reached a maximum at the northern-most stations (Juranek et al. 2020) (Fig. 1E, F). Heterotrophic bacterial cell abundances at the surface also increased with latitude by tenfold across both transects (Fig. 1G).

Surface dissolved iron concentrations showed different latitudinal trends between the two years (Fig. 2A). In April 2016, surface dissolved iron concentrations were highest in the NPSG (0.13-0.31 nM) and generally decreased with increasing latitude, whereas in June 2017, surface dissolved iron concentrations were highest in the NPTZ (0.20-0.51 nM) (Pinedo-González et al. 2020). The peak in dissolved iron concentrations at 35°N in June 2017 was interpreted to be due to atmospheric deposition of anthropogenic aerosols from Asia, based on the isotopic composition of dissolved iron and lead (Pinedo-González et al. 2020). The difference in iron distributions between the two cruises was likely caused by the seasonal supply of aerosol-derived iron to the North Pacific Ocean, as depth-averaged satellite observations of aerosol optical depth (AOD) show higher aerosol concentrations over the region of our cruise transect in spring to early summer (March-July; Supp. Fig. 2).

**Figure 2.**
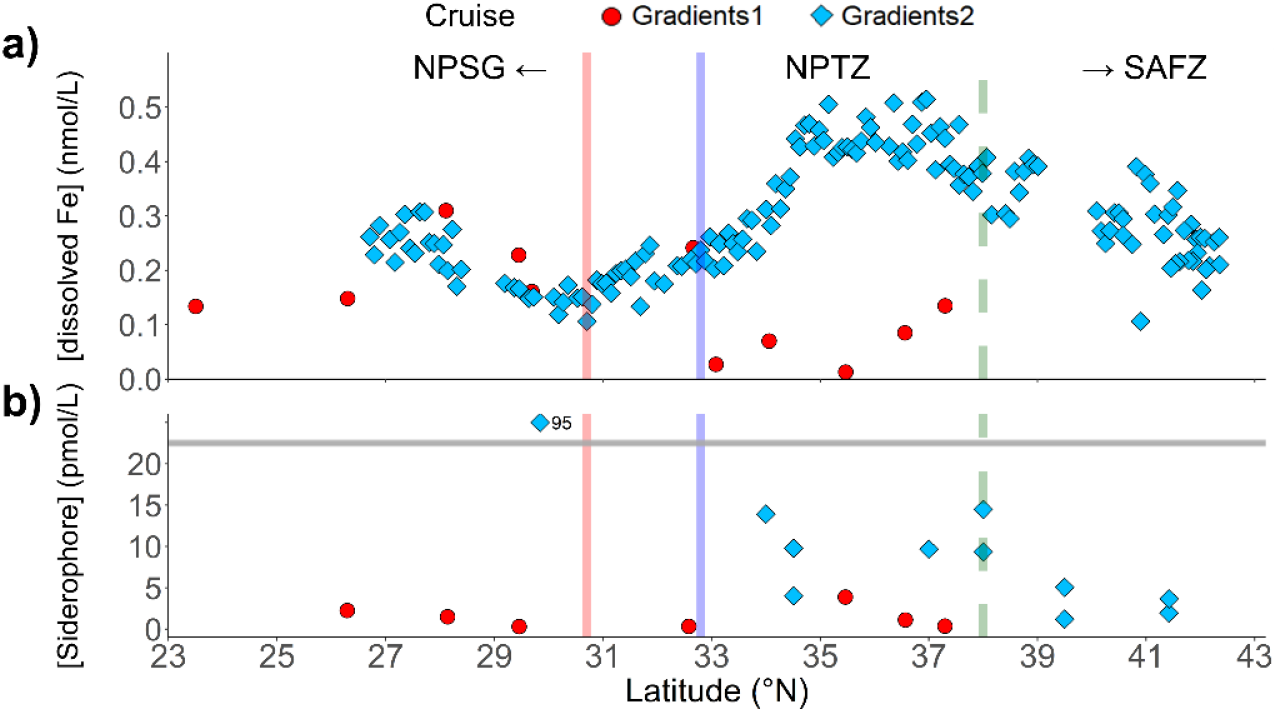
Dissolved iron and siderophore concentrations in surface waters along the two cruise transects. A) Dissolved iron concentrations in surface waters collected with a trace metal clean towfish from Gradients 1 (red) and Gradients 2 (blue). B) Siderophore concentrations in surface waters (0 – 25 m) from Gradients 1 (red) and Gradients 2 (blue). One data point from Gradients 2 had siderophore concentrations outside the y-axis range of the plot (above the horizontal gray y-axis break line), and its concentrations is noted at the right of the data point. Vertical red and blue solid lines indicate location of subtropical fronts from Gradients 1 and 2 cruises respectively; green dashed line indicates location of the subarctic front from Gradients 2.

### Siderophore distributions in the North Pacific

Dissolved siderophore concentrations in surface waters (0 - 25 m) differed between the April 2016 and June 2017 transects (Fig. 2B). In April 2016, surface siderophore concentrations (1.4 ± 1.3 pM) were relatively uniform across the transect, with no statistically significant difference between the NPSG (1.4 ± 1.0 pM) and the NPTZ (1.4 ± 1.7 pM) (Mann-Whitney U test, *p* = 1). In contrast, surface siderophore concentrations in June 2017 were higher and more variable than in 2016, with average concentrations of 14.3 ± 25.9 pM. In addition, siderophore concentrations in the NPTZ (9.3 ± 4.1 pM) were higher than concentrations in the SAFZ (3.0 ± 1.7 pM) (Mann-Whitney U test, *p* = 0.06).

Siderophore concentrations below the surface (25 - 400 m) were not significantly different from surface concentrations in either 2016 or 2017 (Mann-Whitney U test, p = 0.72 and 0.11 respectively). Siderophore concentrations at depth were slightly higher in the NPSG (2.0 ± 1.6 pM) than in the NPTZ (0.9 ± 0.6 pM) (Mann-Whitney U test, *p* = 0.07) in April 2016, while in June 2017 siderophore concentrations did not show a significant difference between the NPTZ (12.8 ± 23.5 pM) and SAFZ (3.3 ± 3.1 pM) (Mann-Whitney U test, *p* = 0.13) (Fig. 3B).

**Figure 3.**
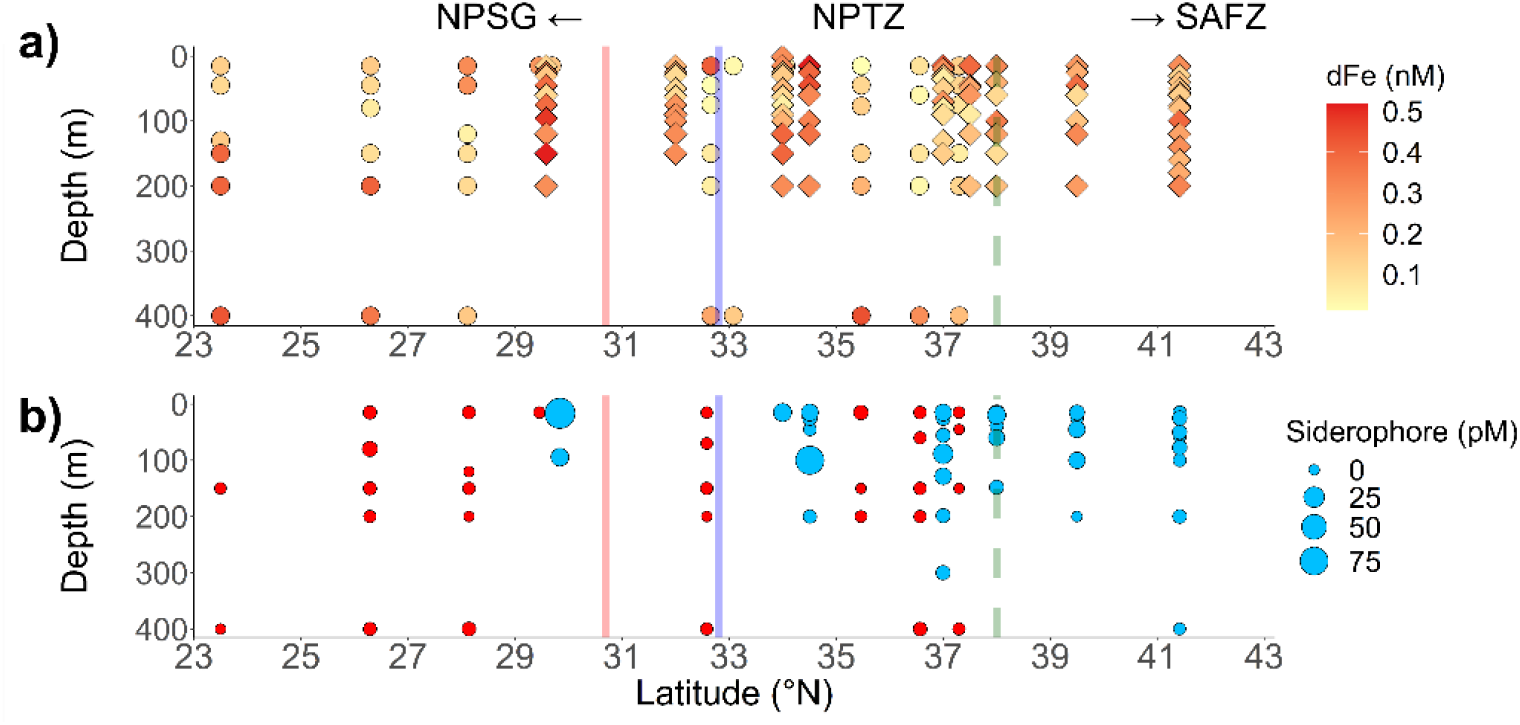
Depth profiles of dissolved iron and siderophore concentrations along the two cruise transects. A) Depth profiles of dissolved iron concentrations from Gradients 1 (circles) and Gradients 2 (diamonds) across 0 – 400 m from CTD casts. B) Depth profiles of dissolved siderophore concentrations from Gradients 1 (red) and Gradients 2 (blue) across 0 – 400 m. Point sizes are proportional to siderophore concentrations. Vertical red and blue solid lines indicate location of subtropical fronts from Gradients 1 and 2 cruises respectively; green dashed line indicates the location of the subarctic front from Gradients 2.

### Siderophore identities in the North Pacific

Among the siderophores we identified with high confidence (levels 1-3) in April 2016 (9 siderophores) and June 2017 (11 siderophores), ferrioxamines were the most common type of siderophore detected and were found in samples from Gradients 1 (Fig. 4) and Gradients 2 (Fig. 5). Ferrioxamine E and G1 were detected in both years (Figs. 4A-B, 5A-B), whereas ferrioxamine B was only detected in April 2016 (Fig. 4C). Most ferrioxamines were complexed to iron, however in 2 out of 37 ferrioxamine identifications, both iron-bound and apo- (iron-free) ferrioxamines were detected, with concentrations of apo-ferrioxamines approximately a tenth of iron-bound ferrioxamines. Different types of ferrioxamines were found across all depths in both the NPSG and the NPTZ, but not in the SAFZ. In addition, multiple types of ferrioxamines were often detected concurrently in a single location. We also detected a suite of amphibactins (D, E, H, T) complexed to iron in June 2017, and these were mostly found in the NPSG and the southern part of the NPTZ (∼35*°*N), except amphibactin E, which was found across the full range of NPTZ (Fig. 5C-F).

**Figure 4.**
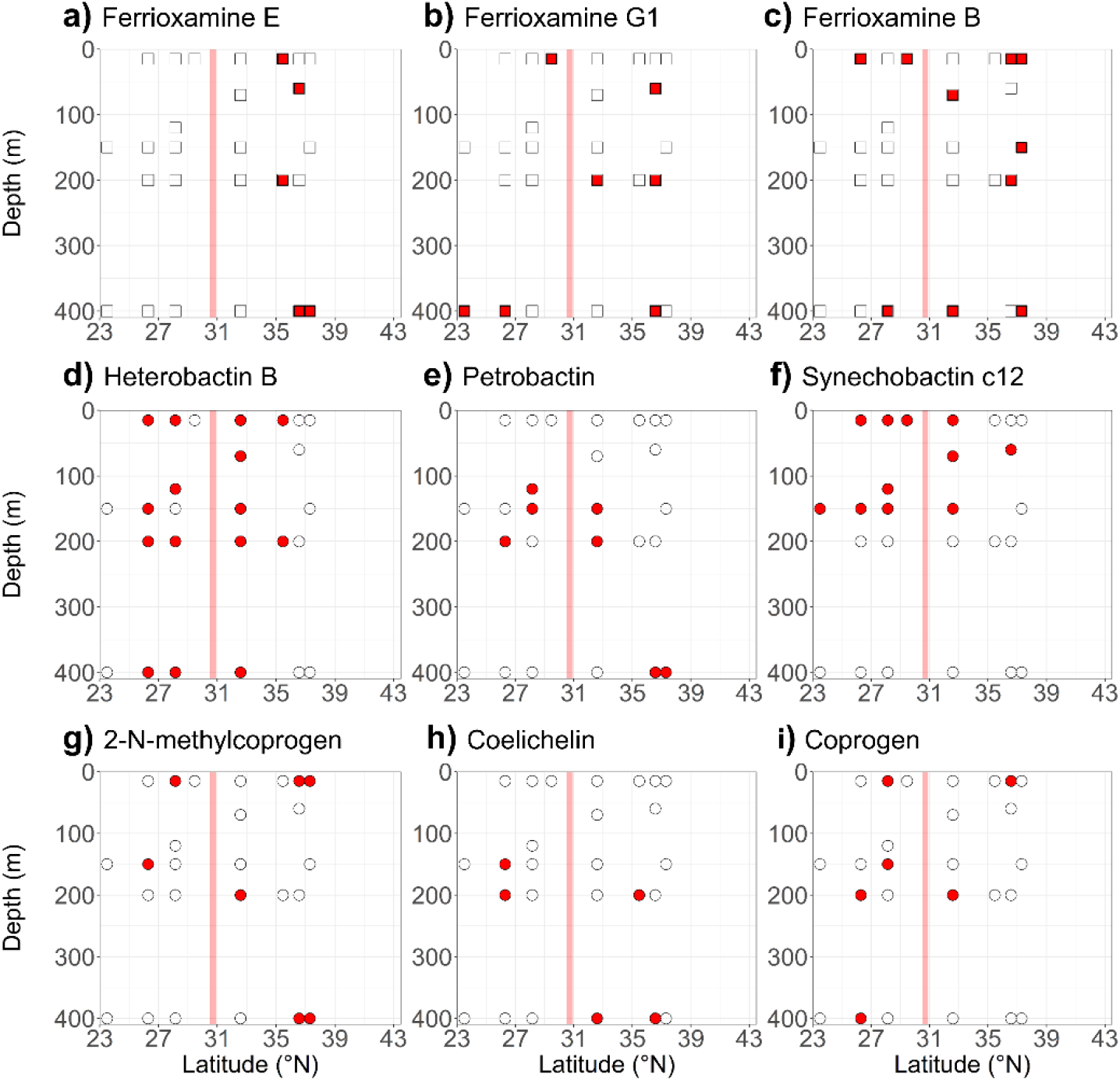
Distributions of siderophores identified with confidence level 1-3 from the Gradients 1 cruise. Red-filled squares (Fe-bound form) and circles (Fe-free form) indicate stations where each siderophore was detected; blank squares and circles show where siderophore was not detected. Vertical red solid line indicates the location of the subtropical front from Gradients 1.

**Figure 5.**
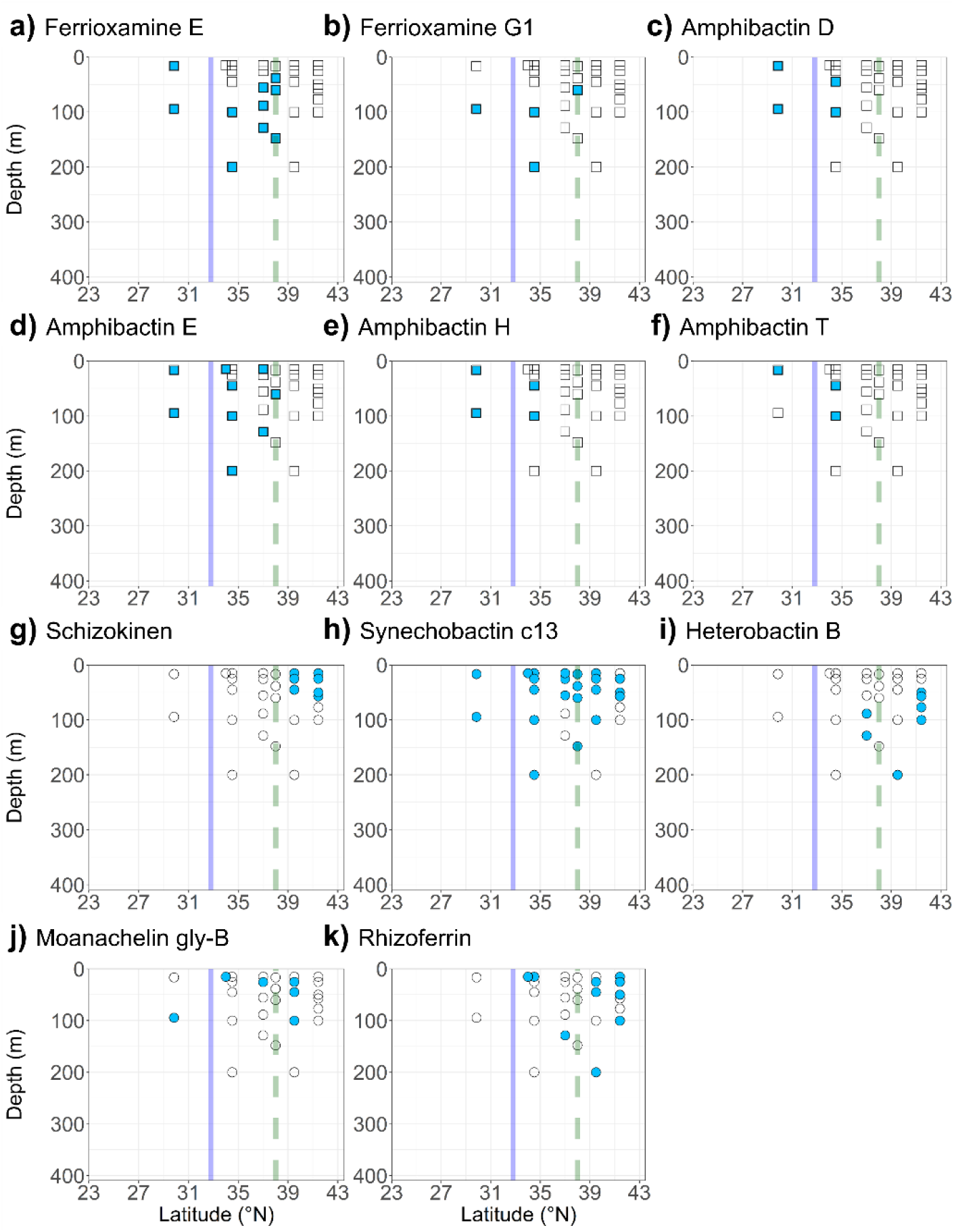
Distributions of siderophores identified with confidence level 1-3 from the Gradients 2 cruise. Blue-filled squares (Fe-bound form) and circles (Fe-free form) indicate stations where each siderophore was detected; blank squares and circles show where siderophore was not detected. Vertical blue solid line and the green dashed line indicate location of the subtropical front and the subarctic front from Gradients 2.

In addition to ferrioxamines and amphibactins, we also detected several photoreactive siderophores in their iron-free forms, based on their MS^1^ and MS^2^ fragmentation data with confidence levels 1-3. Apo forms of synechobactin c12 and c13 were detected consistently across the cruise transects in April 2016 and June 2017, respectively (Fig. 4F, 5H), and were mostly confined to surface waters (0-150 m). Iron-free petrobactin, another photolabile siderophore, was detected in April 2016, at depths between 100 - 400 m (Fig. 4E). North of the subarctic salinity front in June 2017, we also found apo forms of schizokinen (Fig. 5G), another siderophore that may be photoreactive given the α-hydroxy-carboxylic acid group in its structure, although its photoreactivity has not yet been confirmed (Å rstøl and Hohmann-Marriott 2019).

In addition, we detected multiple siderophores not previously detected in the marine environment. Apo forms of heterobactin B were found in both years (Fig. 4C, 5I), although they were most common in the NPSG and the NPTZ in April 2016, while they were found only north of the subarctic salinity front in June 2017. We also found a suite of siderophores known to be produced by marine or terrestrial fungi, including coprogen, 2-N-methylcoprogen and rhizoferrin, sporadically distributed across the transect.

### Connecting genomes and transcriptomes to siderophore measurements

We used FeGenie to explore the genetic potential to use and produce siderophores (Garber et al. 2020) and to search for known siderophore transport and synthesis genes in metagenomic datasets collected from surface waters in both cruises. Known siderophore transport genes were more abundant across latitude based on counts per million mapped reads (CPM: 885 ± 233 and 1158 ± 365; Gradients 1 and 2 respectively) and were an order of magnitude more abundant than known siderophore biosynthesis genes across latitude (CPM: 79 ± 41 and 106 ± 41; Gradients 1 and 2 respectively) (Fig. 6A). Neither the abundances of siderophore transport or biosynthesis genes correlated with latitude during either cruise (R^2^ < 0.1). A positive correlation between the abundances of the two gene sets was observed in both cruises (R^2^ = 0.47 and 0.74 for Gradients 1 and 2 respectively).

**Figure 6.**
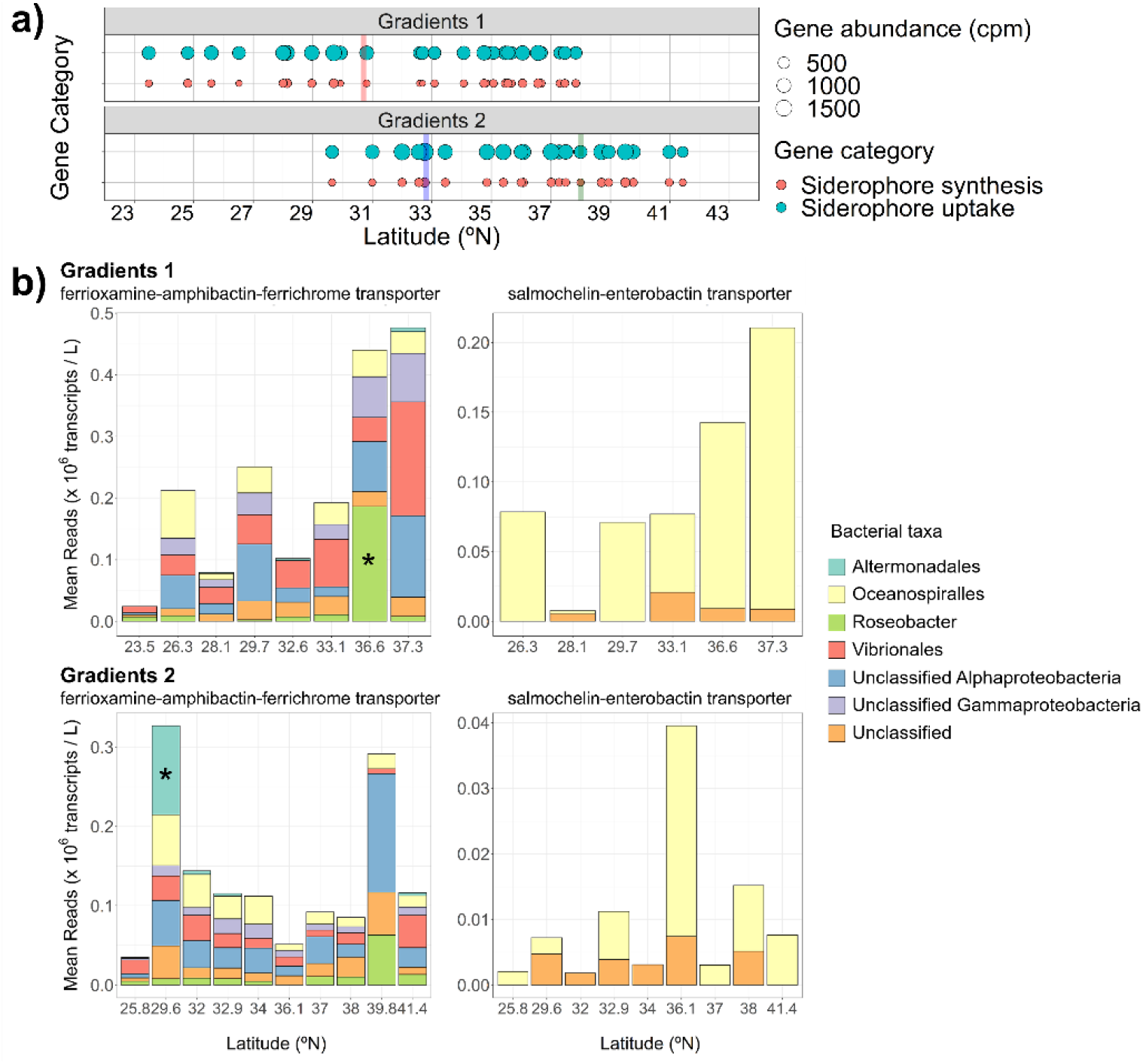
Siderophore uptake genes and transcripts abundances across the two cruise transects. A) Distribution of siderophore uptake (blue) and synthesis (red) genes across latitude at surface waters of Gradients 1 and 2 cruises. Point sizes are proportional to gene abundances (counts per million mapped reads). Vertical red and blue solid lines indicate location of subtropical fronts from Gradients 1 and 2 cruises respectively; green dashed line indicates the location of the subarctic front from Gradients 2. B) Mean abundances of ferrioxamine-amphibactin-ferrichrome and salmochelin-enterobactin uptake transcript homologs (*106 transcripts per liter of seawater) from Gradients 1 and 2 cruises. Taxonomic groups that significantly changed transporter transcription levels at specific latitudes determined by the multiple comparison test (p < 0.05) are marked with stars.

To examine which taxa may be actively cycling siderophores along the cruise transects, we analyzed the transcript levels for homologs of three outer membrane siderophore transporters – ferrioxamine and amphibactin (foxA, desA, fhuA) and salmochelin (fepA, iroN). These outer membrane transporters were chosen as these compounds were detected during both cruises, and outer membrane transporters for these siderophores have been characterized in multiple gammaproteobacterial species. We note that transporter homologs identified in marine microbial reference organisms for ferrioxamine, amphibactin, and ferrichrome, or for salmochelin and enterobactin, could not be distinguished from one another due to their high sequence similarity (Supp. Fig. 1; Supp. Table 1). Here, we report a combined transcript abundance for the transporter homologs, though we also separated transcript inventories for the ferrioxamine-amphibactin-ferrichrome transporter signal based on phylogeny with experimentaly verified sequences (Supp. Fig. 3).

The siderophore transporter homologs were transcribed by taxa belonging predominantly to alpha- and gamma-proteobacteria (Fig. 6B). From both cruises, the alphaproteobacterial Roseobacter group (10 – 13 %) and gammaproteobacterial order Vibrionales (20 – 30 %) contributed most to the total expression of ferrioxamine-amphibactin-ferrichrome transporter transcripts. Mean transcript abundances of ferrioxamine-amphibactin-ferrichrome for transcript homologs differed significantly (one-way ANOVA and post hoc Tukey test) between Roseobacter at 36.6°N in 2016 and Altermonadales at 29.6°N in 2017 and other latitudes (Fig. 6B) (*p* < 0.01). The gammaproteobacterial order Oceanospiralles (29 – 100 %) was the major contributor to total salmochelin-enterobactin transporter transcript abundances (Fig. 6B).

## Discussion

### Siderophore distributions in the North Pacific

Our dataset greatly expands currently available measurements of dissolved siderophore in the ocean. Previous work has been confined to few depth profiles or has focused on surface waters. We found that dissolved siderophores were widely distributed throughout the North Pacific, both in surface waters and throughout the upper mesopelagic (down to 400m), with concentrations comparable to those from earlier studies (0.3 - 20 pM) (Mawji et al. 2008; Boiteau et al. 2016, 2019; Bundy et al. 2018). Two data points at 29.8°N, 16.5 m depth and 37°N, 100 m depth had unusually high concentrations (95.2 pM and 77.7 pM respectively). As all of our dissolved siderophore concentrations are likely representative of a snapshot in time and space, we may have sampled a water mass with a cluster of siderophore-producing microbes or with recently accumulated siderophores from cell death. Nevertheless, dissolved siderophores accounted for 2.4 ± 5.4 % (or 0 – 30%) of the total dissolved iron pool across the North Pacific, suggesting that siderophores are an important component of the iron-binding ligand pool.

Elevated surface siderophore concentrations in the NPTZ in June 2017 may have been caused by higher siderophore production in this area. While a direct correlation between surface siderophore concentrations and heterotrophic bacteria abundance has been observed previously (Mawji et al. 2008), we did not find a strong correlation between surface siderophore concentrations and phytoplankton or bacterial biomass (including NCP, POC, heterotrophic bacteria cell abundances, and chlorophyll concentrations) (Fig. 1E-G, 2B). However, high siderophore concentrations in the NPTZ in June 2017 coincided with a large input of anthropogenic dust to this region (Pinedo-González et al. 2020), and it may be that this external dust input triggered the observed siderophore production. While only a small fraction of dust deposited in the ocean dissolves (approximately 2% but highly variable; Baker and Croot 2010), siderophores can significantly increase the solubility of iron-bearing minerals in dust, such as iron oxides and clay minerals, so that Fe(III) from minerals are released and become available for microbial uptake (Cheah et al. 2003; Kraemer et al. 2005; Parrello et al. 2016). A previous study in the Mediterranean Sea found a positive relationship between the fraction of iron released from dust particles and iron-binding ligand concentrations, with highest dust-derived iron and ligand concentrations in May and June and relative to other months (Wagener et al. 2008). Transient supply of excess iron can result in rapid production and release of siderophores over timescales of days, in order to relieve bacteria from the metabolically costly process of producing siderophores under iron stress (Adly et al. 2015). Therefore, high aeolian dust supply to the North Pacific may have stimulated the microbial communities to increase siderophore production to mobilize and opportunistically take up iron. However, since the dissolved siderophore pool size depends both on siderophore production and release and on siderophore losses through uptake and degradation, different siderophore loss terms across our study sites are also contributing to the observed siderophore concentrations. Estimates of the total siderophore pool size (including the particulate intracellular siderophore pool) or metabolic fluxes of siderophore secretion and uptake are necessary to fully understand how the observed siderophore distribution patterns are being shaped by microbes.

To explore possible environmental drivers of siderophore distributions, including dissolved iron distributions, we further performed a logistic regression analysis using multiple environmental variables to see if any of the variables could successfully predict siderophore distribution. The regression results suggested that latitude was the most effective predictor of siderophore presence, which suggests that siderophore distributions are likely controlled by a combination of physical, chemical and biological factors (see Supplemental Information). Siderophores were also detected below the surface ocean from both years, despite decreased heterotrophic bacterial abundances and increased dissolved iron concentrations with depth (Supp. Fig. 4; Fig. 3A). In the upper mesopelagic waters (150 – 400m), siderophores only make up 0.7 ± 0.8 % (or 0 – 3%) of the total dissolved iron pool. However, previous work found that heterotrophic bacteria from 150 m depth produced siderophores upon addition of *in-situ* sinking particles, suggesting that bacteria are using iron in the sinking particles during regeneration by actively producing and cycling siderophores below the euphotic zone (Bundy et al. 2018). Therefore, while the dissolved siderophore pool size is smaller in the upper mesopelagic relative to the surface waters by up to an order of magnitude, heterotrophic bacteria at depth may more actively cycle siderophores.

### Siderophore identities in the North Pacific

Overall, we observed a wide variety of siderophores with little compositional overlap between the cruises. Ferrioxamines – one of the most commonly detected siderophores in seawater to date – were the only siderophores that were detected on both cruises and in both the NPSG and NPTZ, regions with relatively high iron compared to the SAFZ (Fig. 4A-C, 5A-B). These observations generally agree with previously known distributions of ferrioxamines in iron-replete waters, such as coastal upwelling systems in Southern California and Peru, subtropical North Pacific, and the mid-latitude North Atlantic (Mawji et al. 2008; Boiteau et al. 2016, 2019). The abundances of ferrioxamines in iron-rich waters have been attributed to their extremely strong affinities to iron, which allows bacteria that can utilize ferrioxamines to efficiently compete for iron from dust or sedimentary sources (Mawji et al. 2008; Bundy et al. 2018; Boiteau et al. 2019).

Amphibactins are hydroxamate siderophores with amphiphilic tails that are also frequently detected in seawater, and were found in the NPSG and the southern part of the NPTZ (∼35*°*N) in June 2017 (Fig. 5C-F). Earlier studies detected amphibactins mostly in iron-limited waters such as the eastern tropical Pacific and offshore of the California Current System (Boiteau et al. 2016, 2019). In particular, Boiteau et al. 2016 observed a marked shift from ferrioxamines in the high iron coastal region to amphibactins in the equatorial Pacific HNLC and hypothesized that amphiphilic siderophores, which can partition into the cell membrane, were favored in the HNLC to minimize diffusive losses of the siderophore in this iron-limited region (Boiteau et al. 2016). However, as amphibactins have also been observed in the NPSG (Bundy et al. 2018), their distributions may be driven not only by dissolved iron concentrations, but also by the distribution of various amphibactin-producing and consuming bacterial taxa, including many strains of *Vibrio*, the hydrocarbon-degrading *Alcanivorax* and nitrogen-fixing *Azotobacter* (Martinez et al. 2003; Vraspir et al. 2011; Kem et al. 2014; Zhang et al. 2019).

In addition to iron-bound ferrioxamines and amphibactins, we detected and identified several photoreactive siderophores in the iron-free form. Iron-free forms of synechobactin and schizokinen were found primarily in surface waters (0 – 200 m) (Fig. 4F, 5G-H). As known producers of synechobactin and schizokinen include *Synechococcus* and *Anabaena* species, and as iron-synechobactin complexes can break down within a few hours when exposed to natural sunlight (Å rstøl and Hohmann-Marriott 2019), the presence of synechobactins and schizokinen in surface waters implies active production of photoreactive siderophores in the surface ocean. These siderophores are not routinely detected due to a previous focus on iron-bound compounds. In contrast, the iron-free form of petrobactin was only detected below the surface ocean (100 – 400 m). The absence of iron-free petrobactin in surface waters may be due to photooxidation of the catecholate moieties on petrobactin (opposed to light-resistant hydroxamate moieties on synechobactin and schizokinen) (Barbeau et al. 2003). Photoreactive siderophores have important implications for iron bioavailability, as photolysis of iron-siderophore complexes results in the reduction of Fe(III) to Fe(II), which in turn may become available for marine organisms that cannot take up iron bound to siderophores (Barbeau et al. 2001). Thus, further exploration of photoactive siderophores, especially in the surface ocean, may reveal their importance to marine carbon and iron cycling.

We also putatively identified heterobactin B and some fungi-derived siderophores. Much less is known about the production of these compounds. To date, heterobactins have been shown to be produced by several strains of *Rhodococcus* (Carrano et al. 2001), a genus of gram positive bacteria related to *Mycobacterium*. Little is known about the genetically diverse *Rhodococcus* in the marine environment, but many strains appear capable of degrading hydrocarbons (Sorkhoh et al. 1990; Sharma and Pant 2000). Siderophore production by marine fungi has also not been well-documented. Siderophores produced by fungi characterized to date are mostly hydroxamate siderophores (Holinsworth and Martin 2009). One fungus has been shown to produce rhizoferrin, a carboxylate siderophore (Drechsel et al. 1991; Thieken and Winkelmann 1992). Marine fungi appear to produce higher concentrations of siderophores than their terrestrial counterparts (Baakza et al. 2004), however the importance of these siderophores in oceanic iron cycling remains unclear.

Overall, we observed high variability in the types of siderophores detected between the two cruises, with the exception of ferrioxamines. Differences in siderophore composition between the two cruises may reflect the fast turnover of siderophores, as previous incubation studies suggest that siderophore production can change on daily or weekly timescales (Mohamed and Gledhill 2017; Bundy et al. 2018). Considering the large compositional difference between the two cruises, it is interesting that the suite of ferrioxamines were identified from both cruises. It may be possible that the current siderophore analysis method may be preferentially extracting and isolating ferrioxamines, especially as ferrioxamines are ubiquitously detected from earlier studies (Boiteau et al. 2016, 2019; Bundy et al. 2018). It may be also possible that ferrioxamines are more commonly observed because they are more common siderophores that can be produced and used by a much wider variety of microbes than other siderophores.

### Connecting genomes and transcriptomes to siderophore distributions

Our dissolved siderophore data provides information about the size of the dissolved siderophore pool rather than the metabolic fluxes of siderophores. Thus, a snapshot of siderophore distributions alone may not reflect the iron status of microbes in the marine environment. To gain further insight into the interactions between the dissolved siderophore pool and microbial activity, we used environmental metagenomes to evaluate the genetic potential for siderophore biosynthesis in communities across latitudes, then further evaluated the *in situ* iron status of microbes on short timescales by measuring the transcript levels of iron-siderophore complex transporters.

Siderophore transport and biosynthesis potential was widely represented across the transects. Siderophore transport genes were roughly ten-fold more abundant than biosynthesis genes (Fig. 6A), consistent with the results of previous marine metagenome analyses (Hopkinson and Barbeau 2012; Toulza et al. 2012). As siderophore production is a metabolically expensive process, the scarcity of siderophore biosynthesis genes relative to siderophore transporter genes could be an example of the Black Queen Hypothesis, in which evolutionary selection may have caused most microbes to lose the ability to synthesize siderophores, but left a small subset of the microbial community that retained the ability to produce siderophores and a larger subset that retained the ability to take up siderophore-bound iron (Morris et al. 2012; Mas et al. 2016; Lu et al. 2020).

The relative abundance of siderophore transport and biosynthesis gene abundances did not show clear latitudinal trends across the transects (Fig. 6A). This may reflect that siderophore-related genes are highly conserved across these transects, or more generally, they are common in the open ocean, where surface dissolved iron concentrations are consistently below 0.5 nM without external inputs from rivers, sediments and atmospheric deposition (Moore and Braucher 2007) (Fig. 2A). For example, TonB-dependent outer membrane transporters targeting siderophores as substrates are globally common in open ocean datasets (Tang et al. 2012). Siderophore biosynthesis genes also appear to be largely uniform across different ocean areas, though less frequent in high-iron environments such as hydrothermal vents, rivers and aquifers (Garber et al. 2020). Together, these results suggest that the potential for siderophore production and siderophore-mediated iron uptake is consistently widespread across the open ocean. Therefore, the prevalence of siderophore transport and biosynthesis potential across our cruise transects implies that latitudinal differences in siderophore distribution are not a function of the genetic potential for siderophore uptake and synthesis.

We examined transcript levels of siderophore transporter homologs across the cruise transects to infer when these transporters are in use. We focused on the outer membrane transporters for ferrioxamine, amphibactin, and salmochelin rather than the biosynthesis genes for these siderophores, as these latter genes have not yet been fully characterized.

Putative transporter homologs of these siderophores were predominantly transcribed by Gammaproteobacteria (Fig. 6B), which agrees with earlier studies that repeatedly found members of Gammaproteobacteria to possess a wide range of siderophore transporters relative to other bacterial groups (Hopkinson and Barbeau 2012; Tang et al. 2012; Debeljak et al. 2019). We expected to find amphibactin and ferrioxamine transporter genes transcribed by Gammaproteobacteria, as amphibactins and ferrioxamines are currently known to be produced by multiple gammaproteobacterial taxa, including *Vibrio, Alcanivorax, Azotobacter*, and Actinobacteria (Martinez et al. 2003; Vraspir et al. 2011; Kem et al. 2014; Wang et al. 2014; Zhang et al. 2019). Interestingly, we also detected ferrioxamine-amphibactin-ferrichrome transporter homologs in Alphaproteobacteria reference genomes (Supp. Fig. 1) and detected expression of these homologs in the metatranscriptomes (Fig. 6B). This may indicate that ferrioxamines and amphibactins are used by a wider range of microbes than previously known. No studies to date indicate ferrioxamine and amphibactins can be synthesized or utilized by taxa other than Gammaproteobacteria, in part because they have been the focus of laboratory studies (Hopkinson and Barbeau 2012). However, multiple Rhodobacterales species, including Rhodobacter, Roseovirus, Sulfitobacter and Oceanicola, have genes to transport ferrichrome, a hydroxamate-type siderophore similar to ferrioxamine and amphibactin (Rodionov et al. 2006; Hider and Kong 2010). The high sequence similarity (48 – 100 %) between ferrioxamine, amphibactin and ferrichrome transporters makes it difficult to distinguish them from one another. Thus, some of the transcripts recruited by our reference tree may be ferrichrome uptake transporters, rather than ferrioxamine or amphibactin transporters. Although we did not detect ferrichrome in our study, it is possible that the ferrichrome pool has a rapid turnover and its concentration was too low to be detected. Additional studies are required to discern the specific function(s) of these transporter homologs.

Ferrioxamine and amphibactin transporter transcript inventories from Vibrionales were generally more abundant at northern stations (Fig. 6B, Supp. Fig. 3). Vibrionales are copiotrophic bacteria that are usually present in low abundance, but can quickly grow with additional nutrient pulses from bloom events (Gilbert et al. 2012; Westrich et al. 2016). Multiple species of *Vibrio* can produce and take up amphibactin, and at least one species of Vibrio was capable of exogenous ferrioxamine uptake (Gauglitz et al. 2021). Higher ferrioxamine and amphibactin transporter transcript abundances at higher latitudes by Vibrionales may reflect higher abundances of Vibrionales or increased induction of siderophore-mediated iron uptake pathways in this environment.

Ferrioxamine and amphibactin transporter transcripts from Altermonadales at 29.6°N in 2017 were more abundant relative to other latitudes (Fig. 6B). Ferrioxamine transporter transcripts may be produced by taxa such as Pseudoalteromonas and Alteromonas, which are able to assimilate ferrioxamine-bound iron (Armstrong et al. 2004). However, it is currently unclear whether high transcript abundance at 29.6°N was caused by significant differences in transcription rates or a more abundant Altermonadales population because we cannot normalize the transcript abundances to microbial biomass.

While ferrioxamine and amphibactin transporter transcripts were produced by multiple Gammaproteobacterial and possibly Alphaproteobacterial taxa, transcript homologs of the salmochelin-enterobactin transporter were predominantly detected in the Gammaproteobacterial lineage Oceanospiralles. Salmochelin is a C-glycosylated derivative of enterobactin, and both siderophores are known to be produced by the Enterobacterales, including *Escherichia coli, Salmonella* and *Klebsiella* (Hantke et al. 2003; Müller et al. 2009). While iroN and fepA genes are often annotated as salmochelin and enterobactin transporter genes respectively, both receptors can transport iron-bound salmochelin and enterobactin (Watts et al. 2012), which makes discerning these transport functions from each other not possible. Our observation that salmochelin-enterobactin transporter homologs are restricted primarily to Oceanospiralles, as well as the low abundance of associated transcripts, suggests that salmochelin or enterobactin may be more “specific” siderophores utilized by a smaller subset of microbes than ferrioxamines or amphibactins. Given salmochelin and enterobactin uptake is limited to fewer microbes compared to ferrioxamine and amphibactin uptake, their production may also be low, considering the high metabolic cost of siderophore production. If so, this would result in much lower concentrations of salmochelin and enterobactin in the ocean than ferrioxamines and amphibactins, which may partly explain why we did not find enterobactin in this study (although we putatively identified salmochelin with low confidence). Future characterization of salmochelin and enterobactin biosynthesis pathways may help us address this hypothesis by allowing the analysis of transcripts involved in salmochelin and enterobactin biosynthesis in the environment. Nevertheless, changes in the taxonomic composition of microbial groups that transcribe siderophore transporters suggest that several important bacterial taxa can compete for iron at different latitudes, perhaps contributing to niche differentiation in taxa that actively take up siderophore-bound iron.

## Conclusions

The widespread presence of siderophores across a large range in latitude and depth in the North Pacific highlights the ubiquitous use of siderophores in oceanic iron cycling. Siderophores make up 2.4 ± 5.4 % (or 0 – 30%) of the total dissolved iron pool in our study sites. Siderophore distribution patterns were not directly correlated to any single environmental parameter we measured during the two cruises, such as microbial biomass, dissolved iron or the genetic potential of siderophore production and uptake, but were rather shaped by an interplay of multiple factors, particularly microbial production and uptake. A wide variety of siderophores were identified, including ferrioxamines and amphibactins – the two most commonly identified siderophores in the ocean thus far – as well as putative photoreactive and fungal siderophores. In addition, ferrioxamine, amphibactin and salmochelin were actively used by distinct microbial groups within Gammaproteobacteria and potentially Alphaproteobacteria at different latitudes. Altogether, these results suggest that diverse microbial taxa are primarily driving the patterns of siderophore distributions. While our measurements represent a snapshot of siderophore concentrations at the time of our sampling, a time-series of repeated dissolved siderophore profiles or intracellular siderophore secretion and uptake rate measurements may provide additional insight on the timescale of siderophore production, uptake and recycling by microbes, and will provide key insights into how microbes regulate iron availability in the open ocean.

## Data availability statement

Temperature, salinity, nutrients, dissolved iron, POC, NCP, flow cytometry, total siderophore concentrations and identifications are available at Simons CMAP (https://simonscmap.com/) and Zenodo (http://Zenodo.org). Raw mass spectrometry data files will be deposited to MASSive (https://massive.ucsd.edu/ProteoSAFe/static/massive.jsp) with publication. Raw sequence data for metatranscriptomes from Gradients 1 are available under BioProject PRJNA492143. Raw sequence data for metatranscriptomes from Gradients 2 and metagenomes from Gradients 1 and 2 will be available with publication.

## Acknowledgments

This work was supported by the Simons Foundation (426570 to R.M.B, S.G.J., E.V.A., D.L., L.J., A.E.I. and R.M.B.; 823165 to B.P.D.). We thank the SCOPE-Gradients team for assisting with sample collection, sharing data and providing helpful feedback that improved this work. We thank L. Carlson and K. Heal for help with data acquisition on the Orbitrap, T. Ugrai for technical support on the ICP-MS, L. Pnueli and M. Peleg for their contributions to metagenome sample processing and sequence analysis, S. Curless and R. Foreman for nutrient analysis, J. McMillan, K. Cain and A. Hynes for helping collecting and processing flow cytometry samples, D. Karl and A. White for constructive comments on the manuscript, and the captain and crew of the R/V *Marcus G. Langseth* and R/V *Ka’imikai-O-Kanaloa*.

