## Supplementary Information for "Siderophore production and utilization by microbes in the North Pacific Ocean"

Supplementary Table 1 (.xlsx file):

List of siderophores putatively identified from Gradients 1 and 2 cruises and annotated with confidence levels

Supplementary Table 2 (.xlsx file):

List of ferrioxamine, amphibactin and salmochelin outer membrane transporter protein sequences from reference genomes, and BLASTp search results for homologous ferrioxamine-amphibactin-ferrichrome and salmochelin-enterobactin transporter sequences

Included below:

Supplemental Methods and Results

Supplementary Table 3

Supplementary Figures 1 – 4

### Supplemental Methods and Results

#### *Potential environmental predictors of siderophore distributions*

To gain insight on which environmental parameters could predict the presence of siderophores in each sampling location, we performed a multiple logistic regression analysis on all putatively identified siderophores. The regression models included eight environmental variables (depth, latitude, temperature, salinity, photosynthetically active radiation (PAR), N + N, SRP and dissolved iron concentrations) that were standardized prior to the statistical analysis to account for different scales of measurement. To minimize the effect of multicollinearity between these predictor variables and to reduce the dimensions of the datasets, a principal component analysis (PCA) was first conducted using samples from both cruises. The first three principal components (PCs) accounted for 91% of the variation, so these three PCs were used to build initial multiple logistic regression models. Among 78 putatively identified siderophores (levels 1-4), we only built models built for 21 siderophores that were neither very widely or rarely detected (between 15 – 85% of all sampling locations) to avoid skewing model predictions using too many or too few zeros. For the 21 compounds, each PC in the initial model was substituted for environmental variables that were contributing most strongly to each PC. The best performing models were initially selected based on lowest Akaike information criterion (AIC) values, but to avoid selecting highly correlated predictors, models with predictors that exceeded variance inflation factor (VIF) values of 3 were rejected and the next best models were selected.

Our analysis was limited in that the model parameters evaluated did not include biological parameters such as productivity, biomass and microbial community composition, which are potentially important parameters for siderophore predictions, because discrepancies in sampling locations made it difficult to pair some of our biological data directly with the siderophore measurements. In addition, the accuracy of the current regression models was further limited by both the small size of our dataset and the range of environmental parameters measured during the two cruises. We believe that additional siderophore measurements from different areas of the ocean would likely improve the performance of current models.

Despite these limitations, we find that latitude was the most effective predictor variable among the variables tested in both cruises from our regression models (included in 16 models out of 21) (Supp. Table 3). The regression coefficients of latitude for compounds detected during the April 2016 cruise were mostly negative (8 out of 9), whereas those for compounds detected during the June 2017 cruise were all positive. This trend may be partly due to the differences in sampling locations between the two cruises, since dissolved siderophore samples were collected from relatively lower latitudes in the April 2016 cruise than the June 2017 cruise, and the statistical analyses combined data from both cruises.

Following latitude, salinity was also identified as a significant predictor for 4 compounds, although replacing salinity with latitude returned almost equally efficient regression models, with slightly higher AIC scores (5 – 10) relative to the best models. Dissolved iron concentration was also identified as a potentially important predictor for three siderophores found in April 2016, which were all predicted to be more likely to be found with low iron. Lack of predictive power of dissolved iron towards siderophores may be expected, considering that siderophores have been shown to be present in both iron-limiting and iron-replete areas of the ocean (Mawji et al. 2008; Boiteau et al. 2016, 2019; Bundy et al. 2018).

N + N concentrations and depth were found as a significant predictor in two and one compounds respectively, but the relationship between siderophores and macronutrients or depth are still largely unclear. There have been contradicting reports from culture studies about whether macronutrients can significantly affect siderophore, even within the same genus that produce the same siderophores (Fallahzadeh et al. 2010; Vindeirinho et al. 2021). Previous studies that identified siderophores from seawater have also largely focused on surface waters, which limits our knowledge on siderophores at depth.

Supplementary Table 3. Results of multiple logistic regression analyses on each siderophores

Each row on the table shows individual siderophores, for which one or more variables were identified as significant ($p < 0.05$) predictors in each logistic regression model. The top column shows the environmental variables, and the second column shows whether each siderophore was identified from Gradients 1 (G1) or Gradients 2 (G2) cruises. Positive and negative regression coefficients for each variable are noted in cells filled with red and blue respectively.

| **Siderophore** | **Latitude** | | **Salinity** | | **N + N** | | **dFe** | | **Depth** | |
|---|---|---|---|---|---|---|---|---|---|---|
| | G1 | G2 | G1 | G2 | G1 | G2 | G1 | G2 | G1 | G2 |
| Agrobactin A | | | 2.97 | | | | -6.61 | | | |
| Amonabactin P693 | -0.38 | | | | 0.48 | | -8.31 | | | |
| Carboxymycobactins 108_1 | | 0.76 | | | | | | | | |
| Desferrioxamine T1 | -0.25 | | | | | | | | | |
| Desferrioxamine T2 | -0.26 | | | | | | | | | |
| Desferrioxamine T3 | -0.16 | | | | | | | | | |
| Erythrochelin | | 0.29 | | | | | | | | |
| Heterobactin B | -0.35 | | | | | | | | | |
| Micacocidin | -0.24 | | | | | | | | | |
| Mycobactin H | | 0.66 | | | | | | | | |
| Mycobactin P | | 0.39 | | | | | | | | |
| Mycobactin R | | 0.69 | | | | | | | | |
| Mycobactin T | | | -1.62 | | | | | | | |
| Neocoprogen II | | 0.45 | | | | | | | | |
| Parabactin A | -0.28 | | | | | | | | | |
| Protochelin | -0.24 | | | | | | | | | |
| Salmochelin SX | | | | | | | | | -0.04 | |
| Synechobactin c12 | | | 3.54 | | | | | | | |
| Synechobactin c13 | 0.35 | | | | -0.44 | | | | | |
| Vanchrobactin | | 0.33 | | | | | | | | |
| Vibriobactin | | | 3.32 | | | | -6.77 | | | |

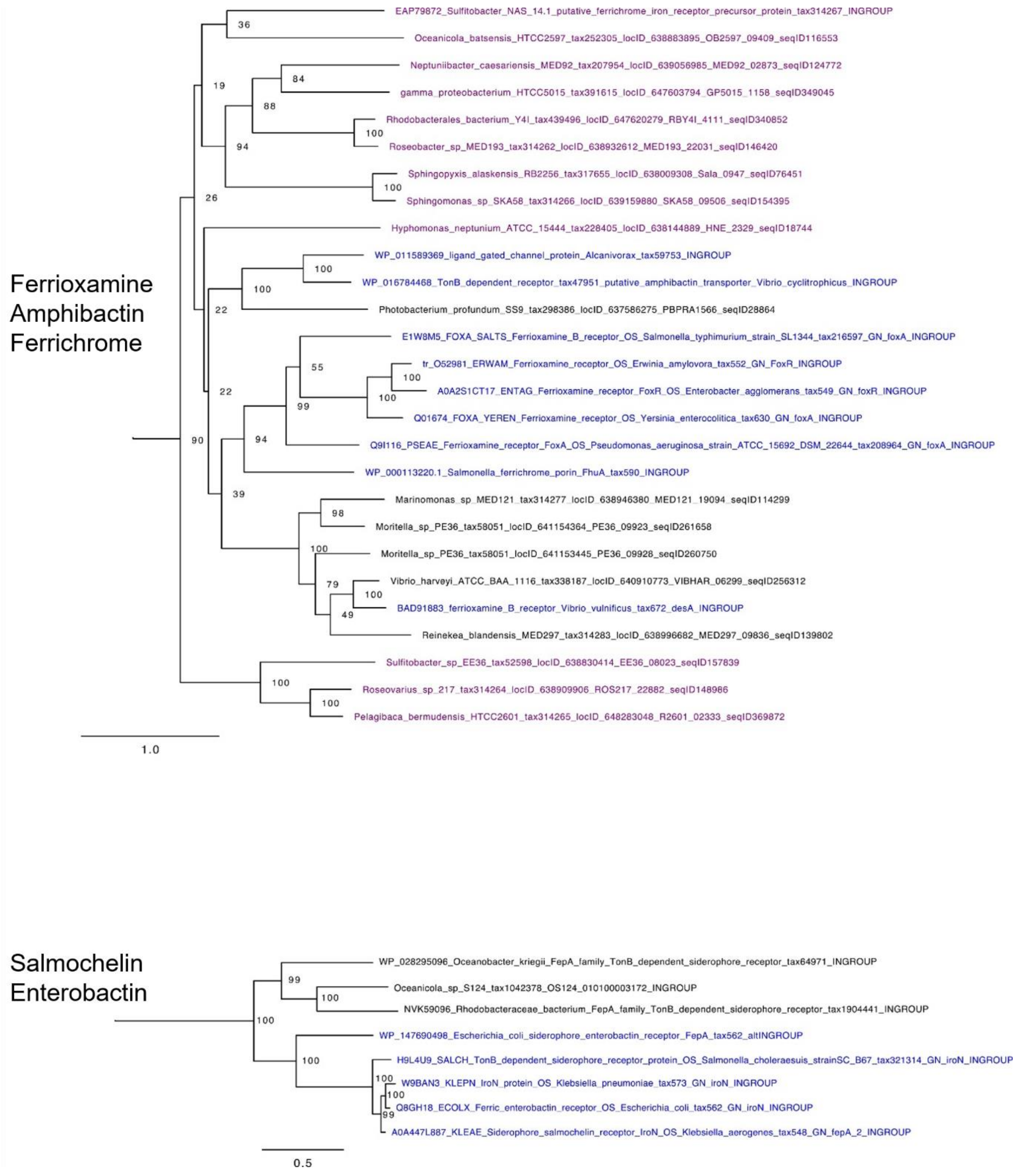


Supplementary Figure 1. Phylogenetic reference trees used to recruit ferrioxamine-amphibactin-ferrichrome transporters and salmochelin-enterobactin transporter homologs. Blue sequences are experimentally verified protein sequences, while black and purple sequences are more and less similar homologs to the experimentally verified protein sequences respectively. Values at the nodes show the number of times the clade appeared in 100 bootstrapped data sets, and the scale bar indicates number of substitutions per site.

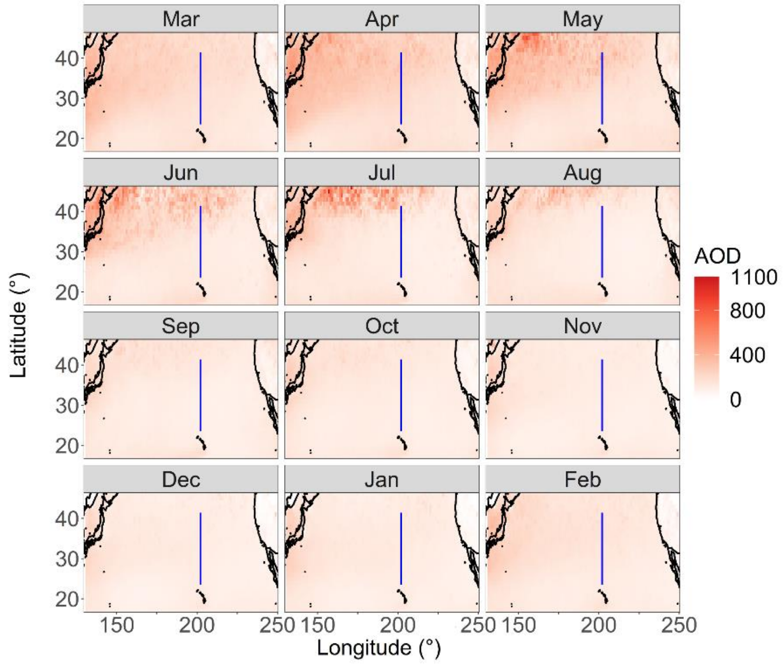


Supplementary Figure 2. Monthly aerosol optical depth (AOD) distribution over the North Pacific Ocean, from NASA's MODIS-Aqua satellite. Monthly data from 2003 to 2018 was averaged over 1 ×1 degree pixels for plotting. Blue line shows both cruise transects from 23.5°N to 41.4°N.

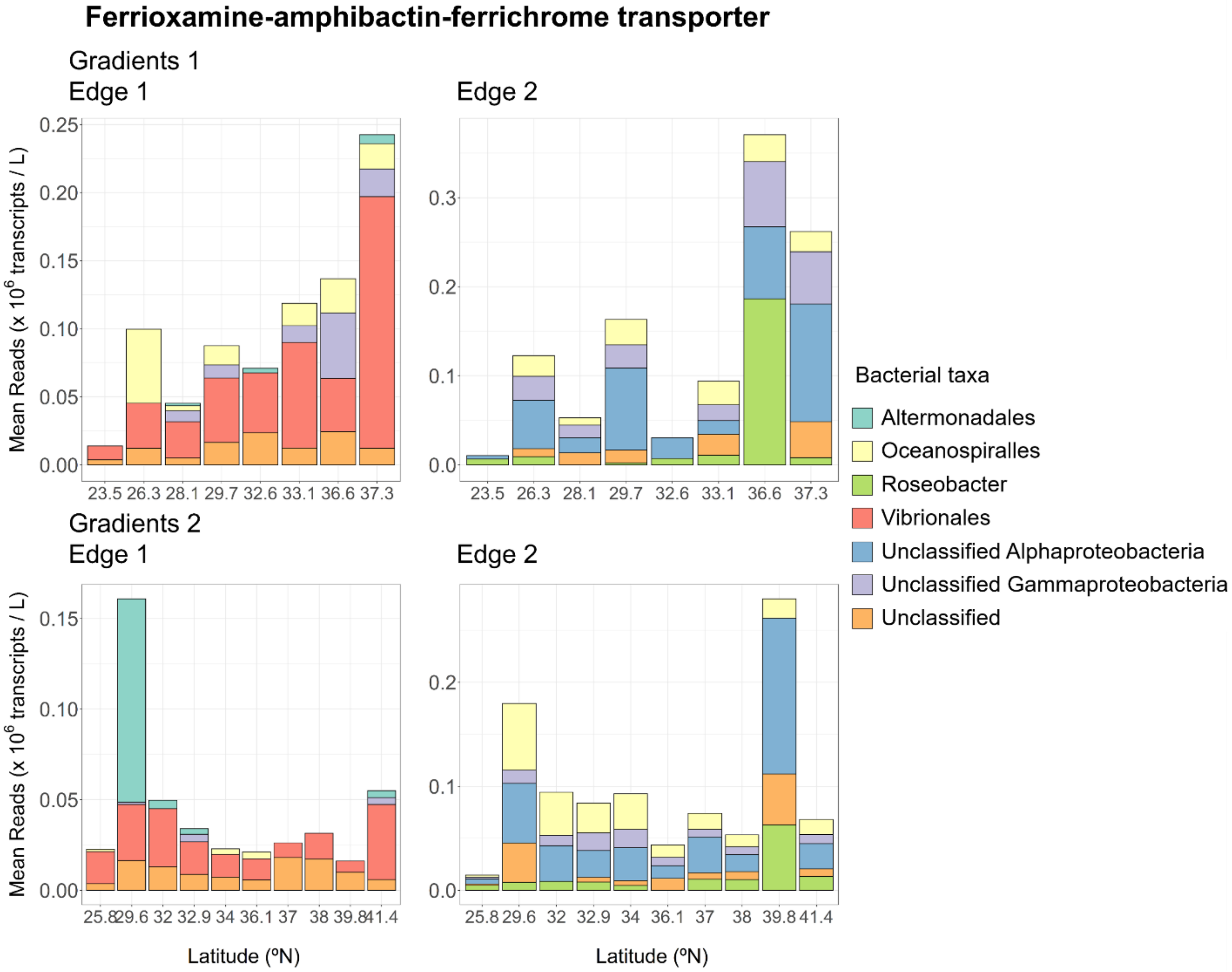


Supplementary Figure 3. Mean abundances of ferrioxamine-amphibactin-ferrichrome uptake transcript homologs (*$10^6$ transcripts per liter of seawater) from Gradients 1 and 2 cruises. Sequences that are more closely related to experimentally verified sequences (black sequences in Supp. Fig. 1) and less closely related (purple sequences in Supp. Fig. 1) were counted separately as Edge 1 and Edge 2.

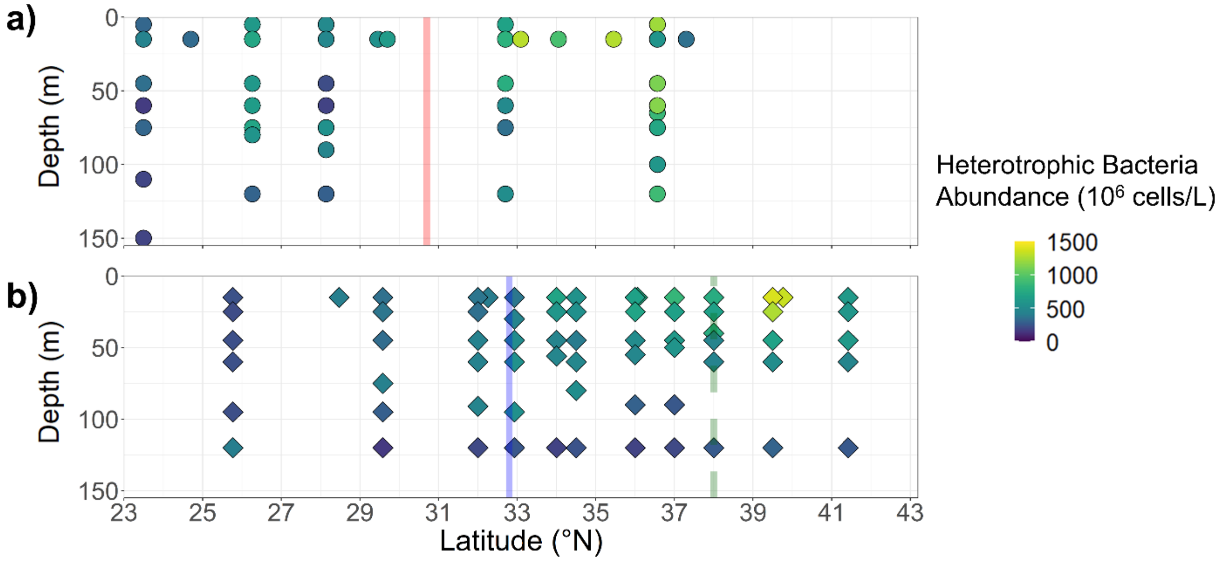


Supplementary Figure 4. Abundance of heterotrophic bacteria determined by flow cytometry from A) Gradients 1 and B) Gradients 2 cruises. Vertical red and blue solid lines indicate the location of subtropical fronts from Gradients 1 and 2 cruises respectively; vertical green dashed line indicates the location of the subarctic front from Gradients 2.
